# Expression of a miRNA targeting mutated SOD1 in astrocytes induces motoneuron plasticity and improves neuromuscular function in ALS mice

**DOI:** 10.1101/2021.01.08.425706

**Authors:** C. Rochat, N. Bernard-Marissal, S. Pradervand, F.E. Perrin, C. Raoul, P. Aebischer, B.L. Schneider

**Author notes:** Current address of corresponding author: Bernard Schneider, EPFL SV PTECH PTBTG, Ch. des Mines 9, CH-1202 Genève, Switzerland. Study was performed in Lausanne, Switzerland.

## Abstract

In amyotrophic lateral sclerosis (ALS) caused by *SOD1* gene mutations, both cell-autonomous and non-cell-autonomous mechanisms lead to the selective degeneration of motoneurons. Here, we evaluate the therapeutic potential of gene therapy targeting mutated *SOD1* in mature astrocytes using mice expressing the mutated SOD1^G93A^ protein. An AAV-gfaABC_1_D vector encoding an artificial microRNA is used to deliver RNA interference against mutated SOD1 selectively in astrocytes. The treatment leads to the progressive rescue of neuromuscular junction occupancy, to the recovery of the compound muscle action potential in the *gastrocnemius* muscle, and significantly improves neuromuscular function. In the spinal cord, gene therapy targeting astrocytes protects a small pool of fast-fatigable motoneurons until disease end stage. In the *gastrocnemius* muscle of the treated *SOD1^G93A^* mice, the fast-twitch type IIb muscle fibers are preserved from atrophy. Axon collateral sprouting is observed together with muscle fiber type grouping indicative of denervation/re-innervation events. The transcriptome profiling of spinal cord motoneurons shows changes in the expression levels of factors regulating the dynamics of microtubules. Gene therapy delivering RNA interference against mutated SOD1 in astrocytes provides therapeutic effects enhancing motoneuron plasticity and improving neuromuscular function in ALS mice.

## Introduction

Amyotrophic Lateral Sclerosis (ALS) is a fatal neurodegenerative disease characterized by the progressive and selective loss of motoneurons (MN) in the cortex, brainstem and spinal cord. Whereas 90% of ALS cases are sporadic, the remaining 10% are familial (fALS). Pathogenic mutations in the gene encoding Cu/Zn superoxide dismutase (*SOD1*) are considered to cause 20% of fALS cases ^1^.

Mice overexpressing mutated SOD1 (mSOD1) replicate the main features of ALS ^2^. In the spinal cord, neurodegeneration follows a specific pattern, characterized by the higher vulnerability of the fast-fatigable MN, which innervate fast-twitch type IIb muscle fibers ^3^. Importantly, non-cell-autonomous pathogenic mechanisms are also involved in MN degeneration ^4–6^. Mutated SOD1 has key pathogenic effects in glial cells including astrocytes, microglial cells and oligodendrocytes. Suppressing mSOD1 in these cell types modifies onset and/or progression of the disease ^5,7,8^.

Diseased astrocytes have major effects on MN, both *in vitro* and *in vivo*. In co-culture systems, astrocytes derived from ALS patients or from ALS mouse models show toxic activities on MN ^9–11^. Furthermore, infusion of culture medium conditioned by primary mouse astrocytes expressing mSOD1 is sufficient to induce MN degeneration and neuromuscular dysfunction in healthy rats ^12^. Although the molecular cause of this toxicity remains poorly understood, some mechanisms have been proposed. Astrocytes from ALS patients and mSOD1 mice have reduced expression of *EAAT2*, which may decrease their ability to properly uptake glutamate, causing excitotoxicity in MN ^13^. Furthermore, ALS may affect the ability of astrocytes to provide essential metabolic and trophic support to MN, for instance by lowering lactate efflux ^14^. Because of an abnormal ratio between pro- and mature nerve growth factor in astrocytes, the p75 signaling pathway was aberrantly up-regulated in MN ^14^. The release of transforming growth factor β1 (TGF-β1) is up-regulated in ALS astrocytes, with possible effects on the inflammatory response and MN survival ^15,16^. Diseased astrocytes can also produce factors that are directly toxic to MN, such as high levels of NO and interferon gamma (IFNγ), which may increase oxidative stress and cause MN death ^9,17,18^. Reactive astrocytes have also been shown to release lipocalin 2 (lcn2), which is a potent mediator of neuronal toxicity ^19^.

Various strategies are explored to prevent the toxic effects of diseased astrocytes in ALS, including RNA interference to lower the expression level of mSOD1 protein ^20–24^. We previously designed an AAV9 vector combined with the gfaABC_1_D promoter driving expression of an artificial microRNA to knockdown human SOD1 (miR SOD1) in astrocytes ^25^. Following intracerebroventricular (ICV) injection in neonatal *SOD1^G93A^* mice, this vector was found to improve the neuromuscular function and prolong animal survival.

Here, we used this approach in *SOD1^G93A^* mice to explore the neuroprotective effects of targeting astrocytes for gene therapy against mSOD1 mediated by RNA interference (RNAi). Towards disease late stage, silencing of mSOD1 led to a significant protection of the motor function, improving mouse performance in specific behavioral tests for muscle strength and motor coordination. Treated mice displayed a partial protection of the vulnerable fast-fatigable MN in the lumbar spinal cord. However, AAV-mediated targeting of astrocytes for expression of miR SOD1 had most significant effects on the occupancy of the neuromuscular junctions (NMJ), which remained highly protected from ALS-induced denervation. In the *gastrocnemius* muscle, gene therapy induced a significant protection of the fast-twitch type IIb muscle fibers. Protection of the neuromuscular junctions was also revealed by events of axonal sprouting and muscle fiber clustering. To further analyze the effects of treated astrocytes on spinal cord MN and identify potential gene candidates implicated in the therapeutic response, we performed a transcriptomic analysis of MN exposed to astrocytes expressing miR SOD1. Altogether, our results show that AAV-mediated gene therapy targeting mSOD1 in astrocytes has clear effects on spinal cord MN in *SOD1^G93A^* mice by promoting functional re-innervation of the skeletal muscle.

## Results

### Expression of miR SOD1 in astrocytes rescues neuromuscular function in *SOD1^G93A^* mice

We assessed the effects of AAV-mediated miR SOD1 gene therapy on disease progression in *SOD1^G93A^* mice. Mice were ICV injected at P2 with the AAV9-gfaABC_1_D:GFP:miR SOD1 vector (AAV-miR SOD1) which drives expression of an artificial anti-human SOD1 miR in astrocytes ^25^. As previously shown ^26^, AAV9 ICV injection in neonatal mice lead to expression of GFP along the spinal cord, including the lumbar region, with similar expression levels at 65 days of age and end stage (day 153-185) (Supplemental Fig. S1). To assess neuroprotective effects of the astrocyte-specific silencing of SOD1, the treated mice were compared to *SOD1^G93A^* mice injected with a similar vector encoding a scramble miR sequence (AAV-miR ctrl), non-injected *SOD1^G93A^* mice (untreated), and wild-type (WT) littermates. Neuromuscular function was assessed along the course of the disease using electromyography and behavioral tests (Fig. 1a). Amplitude of the muscle evoked response (CMAP) was measured every week in the *triceps surae* (Fig. 1b). Until day 73, there were no significant effects of AAV-miR SOD1, as CMAP values declined to 62 ± 4.3 mV in the treated mice, a value only slightly higher than in both control groups of *SOD1^G93A^* mice. CMAP amplitude then increased in the AAV-miR SOD1 group, to reach 88 ± 7.3 mV at day 87, and then stayed stable until day 136. In contrast, CMAP values further decreased in the AAV-miR ctrl and untreated mice. Another miR SOD1-b sequence also targeting human SOD1 showed similar effects, which are therefore not specific to the miR sequence (Supplemental Fig. S2). Muscle strength and motor coordination were evaluated with the grid and rotarod tests, respectively (Fig. 1c, d). Control *SOD1^G93A^* mice showed a progressive loss of muscle strength detectable from day 93 on (Fig. 1c). As compared to control groups, AAV-miR SOD1 significantly improved muscle strength from day 119 until end stage (Fig. 1c). In the rotarod test, the latency to fall was only marginally prolonged in the AAV-miR SOD1 treated mice, when compared to untreated controls, reaching significance only at later stages of the disease (Fig. 1d). These results indicate that silencing of mSOD1 in astrocytes promotes recovery of the neuromuscular function in *SOD1^G93A^* mice, mainly improving muscle strength.

**Figure 1.**
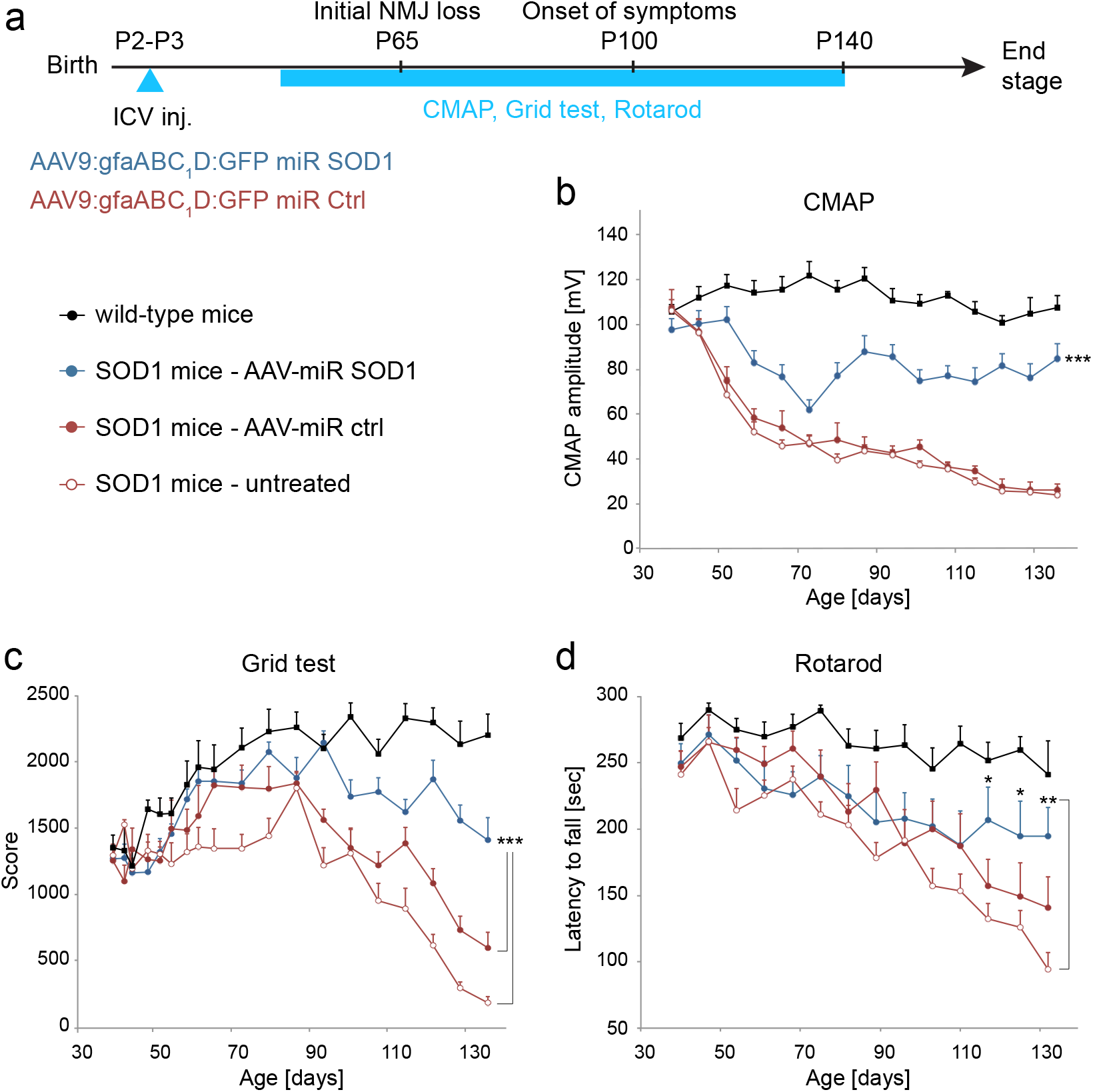
AAV-miR SOD1 targeting of astrocytes preserves neuromuscular function. Neuromuscular function is assessed by electromyography and behavioral testing. (**a**) Schema of the longitudinal experiment indicating the analysis time points. (**b**) Amplitude of the evoked compound muscle action potential (CMAP) recorded in the *triceps surae*. Note the rapid decrease of CMAP amplitude in the untreated and AAV-miR ctrl injected *SOD1^G93A^* mice between day 45 and 66. There is a progressive rescue of CMAP values in the AAV-miR SOD1 treated group, from day 73 onwards. (**c**) The grid test is used to evaluate the strength of the four limbs. Note the marked decrease of the score in the untreated and AAV-miR ctrl injected *SOD1^G93A^* mice, from day 86 on. A significant rescue of muscle strength is observed in the AAV-miR SOD1-treated mice. Statistical analysis for b and c: two-way ANOVA (group x time) repeated measures with Bonferroni *post-hoc* test; \*\*\**P* < 0.001. (**d**) Motor coordination is measured in the Rotarod test. Note the progressive loss of performance in ALS mice, starting at day 75. AAV-miR SOD1 induces a late improvement of the motor coordination, from day 117 on. Statistical analysis: one-way ANOVA with Newman-Keuls post-hoc test; \**P* < 0.05, \*\**P* < 0.01. Data represent mean ± SEM. *n* = 12 mice per group.

### Expression of miR SOD1 in astrocytes protects MN in the lumbar spinal cord but has no effect on the inflammatory response

Next, we analyzed spinal cord tissue at various time points over the disease process, from day 65 (initial drop of the CMAP amplitude) until end stage. The number of choline acetyltransferase (ChAT)-positive MN was determined in the lumbar region of the spinal cord (Fig. 2a, b). At day 65, MN loss was only marginal in *SOD1^G93A^* mice compared to WT mice (Fig. 2b). However, the number of ChAT-positive MN was significantly decreased in all groups at day 140. At end stage, the number of surviving MN reached 10.1 ± 2.1 and 12.2 ± 2 MN (mean ± SEM) per section in the AAV-miR ctrl injected and control untreated ALS mice, respectively. In the AAV-miR SOD1 treated mice however, the number of ChAT-positive MN was significantly increased (14.7 ± 2.1 MN per section) (Fig. 2b).

**Figure 2.**
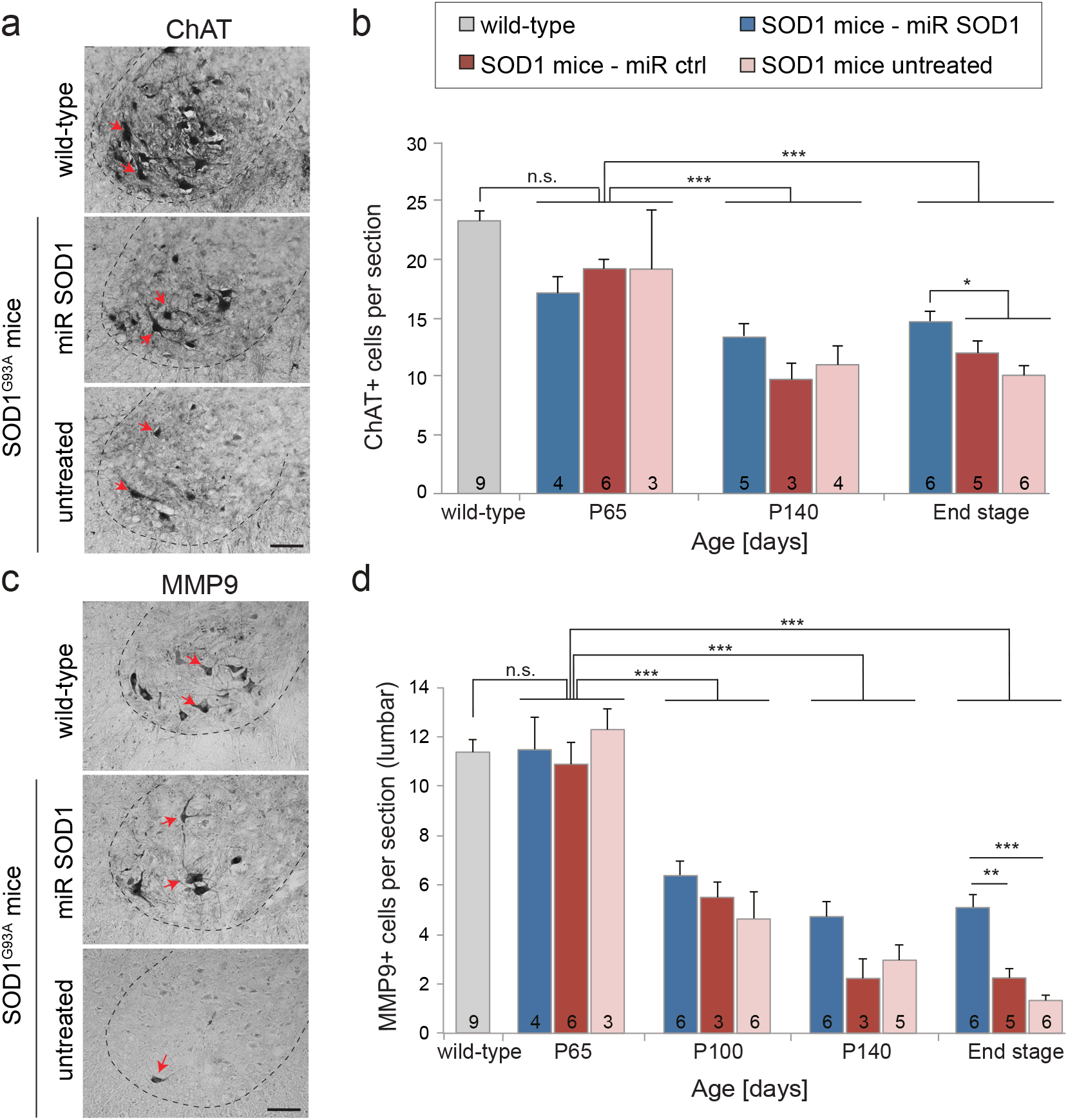
AAV-miR SOD1 targeting of astrocytes has protective effects on fast-fatigable motoneurons in the lumbar spinal cord. (**a**) Representative pictures of ChAT+ MN (red arrows) in the lumbar spinal cord at end stage. (**b**) Quantification of the number of ChAT+ MN per section of the lumbar spinal cord, from 65 days of age until end stage. Note the significant MN protection in the AAV-miR SOD1 group at end stage. For comparison, the grey bar shows the average number of ChAT+ MN per section in adult WT mice (older than 140 days). (**c**) Representative pictures of FF MN highly immunoreactive for MMP9 (white arrows) at end stage. (**d**) Average number of FF MN per section identified by high MMP9 immunoreactivity. There is a significant loss of MMP9+ MN per section in *SOD1^G93A^* mice from day 100 on. Note the significant protection of MMP9+ MN in the AAV-miR SOD1 treated mice at end stage. For comparison, the grey bar shows the average number of MMP9+ MN per section in adult WT mice (older than 65 days). Data represent mean ± SEM. The numbers of replicates per group are indicated in each bar. Statistical analysis: one-way ANOVA with Bonferroni *post-hoc* test; \**P* < 0.05, \*\**P* < 0.01, \*\*\**P* < 0.001. Scale bars: 50 μm.

To assess if AAV-miR SOD1 could prevent astrocytic and microglial activation in the spinal cord, we measured the area of GFAP immunoreactivity and the number of Iba1-positive microglial cells on lumbar sections at day 65, 100 and 140 (Supplemental Fig. S3a-d). As expected, a significant astrocytic and microglial activation was observed in *SOD1^G93A^* mice, from day 100 onwards. However, there were no significant differences in the group treated with AAV-miR SOD1 as compared to control ALS mice. These results indicate that the treatment did not have any major effects on the progression of astrogliosis and microgliosis in *SOD1^G93A^* mice.

### AAV-miR SOD1 injection protects a subpopulation of MMP9 expressing MN

Since all MN subtypes are not equally affected in ALS mice, we further evaluated the effects of the treatment based on the expression of MMP9, a marker highly expressed in fast-fatigable MN, which display highest vulnerability to mSOD1-induced toxicity (Fig. 2c) ^3,27^. At day 65, the number of lumbar MN highly positive for MMP9 was still very similar to WT mice in all groups of *SOD1^G93A^* mice (Fig. 2d). At day 100 however, the loss of fast-fatigable MN already reached 50% and further declined until end stage, at which time point only 1.5 ± 0.5 and 2.2 ± 0.8 fast-fatigable MN were found per lumbar section in the untreated and AAV-miR ctrl injected mice, respectively. In the AAV-miR SOD1 treated mice, the number of MMP9-positive MN remained stable from day 100 onwards. At end stage, we measured 5.1 ± 1.3 MN per lumbar section (mean ± SEM), a number significantly higher than in control groups of *SOD1^G93A^* mice (Fig. 2d). These results show that silencing mSOD1 in astrocytes rescues a subpopulation of fast-fatigable MN with high MMP9 immunoreactivity.

### Expression of miR SOD1 in astrocytes rescues neuromuscular junctions

Despite the partial rescue of spinal cord MN, AAV-miR SOD1 injection induced an effective recovery of CMAP values (see Fig. 1b), which indicates that the treatment may have additional protective effects at the level of the NMJ. To address this possibility, we analyzed NMJ occupancy in the *gastrocnemius* muscle at day 65, 100, 140 and end stage. Acetylcholine receptors were stained with alpha-bungarotoxin to reveal motor endplates, and colocalization with the anti-synaptic vesicle protein 2 (SV2) marker was used to quantify NMJ occupancy (Fig. 3a). Complete overlap between the α-bungarotoxin and SV2 staining accounted for the presence of a fully innervated NMJ (Fig. 3a, b). At day 65, all groups of *SOD1^G93A^* mice, including the group treated with AAV-miR SOD1, displayed a significant loss of fully innervated NMJ as compared to WT mice (AAV-miR SOD1: 61 ± 7% occupancy; WT: 94 ± 2%, mean ± SEM) (Fig. 3b). The innervation of motor endplates further declined until end stage in ALS mice either untreated (13 ± 7%) or injected with AAV-miR ctrl (16 ± 5%). In the AAV-miR SOD1 treated group however, the proportion of fully innervated NMJ increased to 84 ± 5% by 100 days, and remained significantly rescued until end stage (73 ± 7%), only marginally decreased in comparison to WT mice (94 ± 32%) (Fig. 3b). Therefore, injection of the AAV-miR SOD1 vector targeting astrocytes has strong neuroprotective effects on NMJ occupancy.

**Figure 3.**
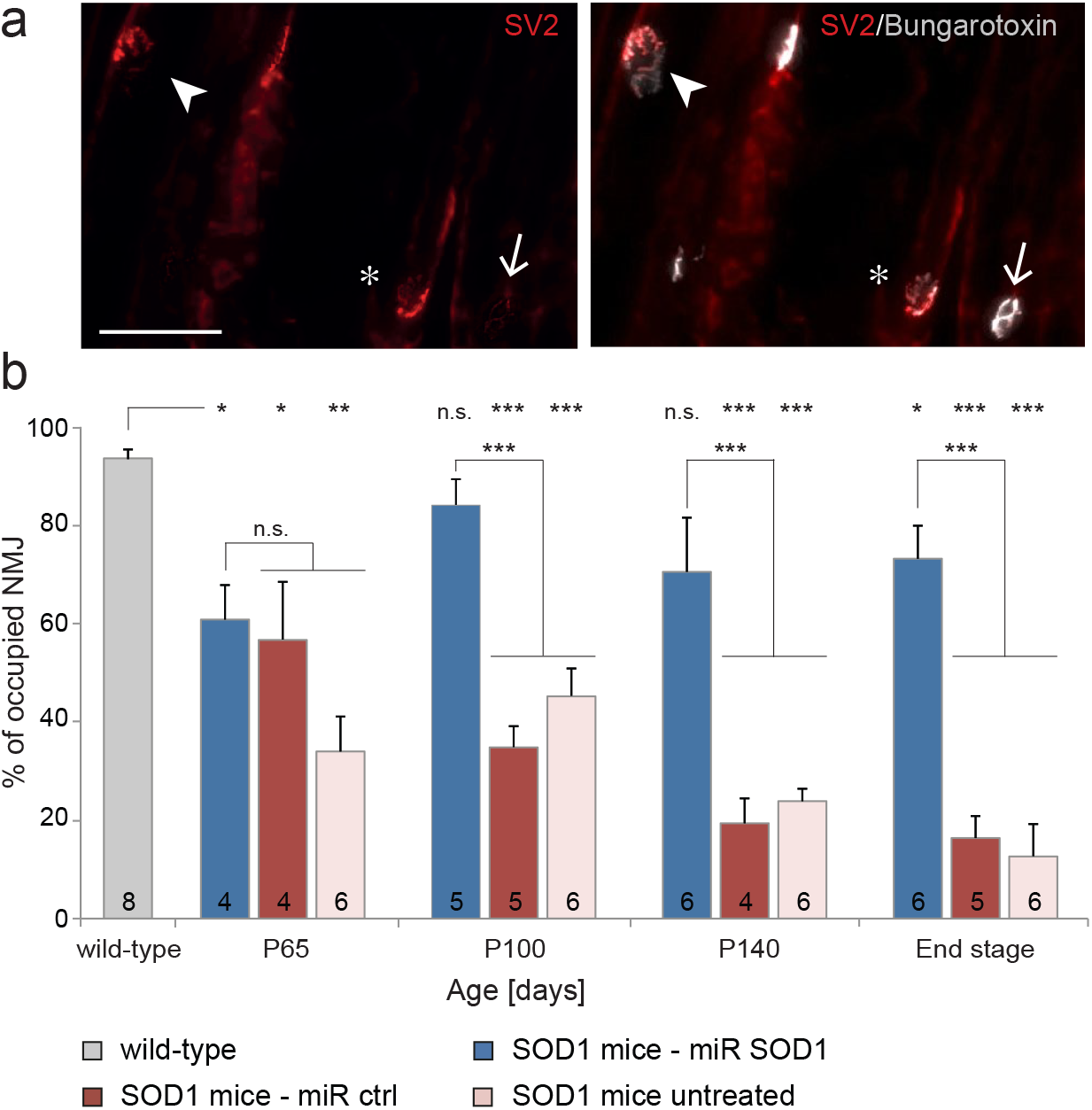
The occupancy of the neuromuscular junctions is rescued in the *gastrocnemius* muscle of *SOD1^G93A^* mice treated with AAV-miR SOD1. Analysis of the innervation of motor endplates in the *gastrocnemius* muscle. (**a**) Immunostaining for SV2 (synaptic marker) and α-bungarotoxin labeling of the motor endplate showing a fully innervated NMJ (*, markers are fully colocalized), a partially innervated NMJ (arrowhead) and an unoccupied motor endplate (arrow). (**b**) Quantification of the percentage of fully innervated NMJ in the *gastrocnemius* muscle. At day 65, NMJ occupancy is significantly decreased in all groups of *SOD1^G93A^* mice. Note the significant rescue of NMJ occupancy at days 100, 140 and end stage in the *gastrocnemius* muscle of AAV-miR SOD1-treated *SOD1^G93A^* mice, as compared to the continuous decrease observed in the AAV-miR ctrl-injected and untreated mice. Data represent mean ± SEM. The number of replicates is indicated in each bar. The grey bar shows the average NMJ occupancy in adult WT mice (older than P65). Statistical analysis: one-way ANOVA with Bonferroni post-hoc test; \**P* < 0.05, \*\**P* < 0.01, \*\*\**P* < 0.001. Scale bar 75 μm.

### Expression of miR SOD1 in astrocytes protects mainly type IIB muscle fibers from atrophy

To further assess the rescue of motor units by AAV-miR SOD1 injection, we analyzed the morphology and composition of the *triceps surae* at end stage. In particular, changes in the number, size and type of muscle fibers were assessed by immunohistochemistry in the *gastrocnemius* and *plantaris* muscles at end stage. In mice, fast-twitch glycolytic type IIB muscle fibers are innervated by fast-fatigable MN, fast-twitch oxidative type IIA fibers by fatigue-resistant MN, and slow-twitch oxidative type I fibers by slow MN. Some fibers display a phenotype which is intermediate between type IIA and IIB, defined as type IIX. Major fiber types were identified using specific staining for myosin heavy chain (MyHC) isoforms (Fig. 4a). Dystrophin staining was used to delineate fiber circumference and quantify the number and mean area of individual muscle fibers (Fig. 4b). As expected, there was a marked muscle atrophy in *SOD1^G93A^* mice either untreated or injected with the AAV-miR ctrl vector. Atrophy was most evident in the *gastrocnemius* muscle, characterized by a major loss of type IIB muscle fibers, but was less pronounced in the *plantaris* (mixed fiber types) and *soleus* (type I and IIA) muscles (Fig. 4a). Compared to WT mice, the total number of fibers in the *gastrocnemius* muscle was decreased by more than 50% in control *SOD1^G93A^* mice (Fig. 4c). In contrast, there was a clear protection of muscle fibers in the AAV-miR SOD1 treated mice. Both the number and the area of the type IIB fibers remained nearly unchanged in the AAV-miR SOD1 treated mice as compared to WT animals, whereas these parameters were dramatically decreased in the control groups (Fig. 4d, e). In contrast, there was no significant difference neither in the number, nor in the area of type IIA, type I and type IIX fibers across groups (Fig. 4d, e). Overall, these results indicate that AAV-miR SOD1 treatment has major protective effects against atrophy of type IIB muscle fibers, consistent with the observed protection of the fast-twitch fatigable MN in the lumbar spinal cord (Fig. 2d).

**Figure 4.**
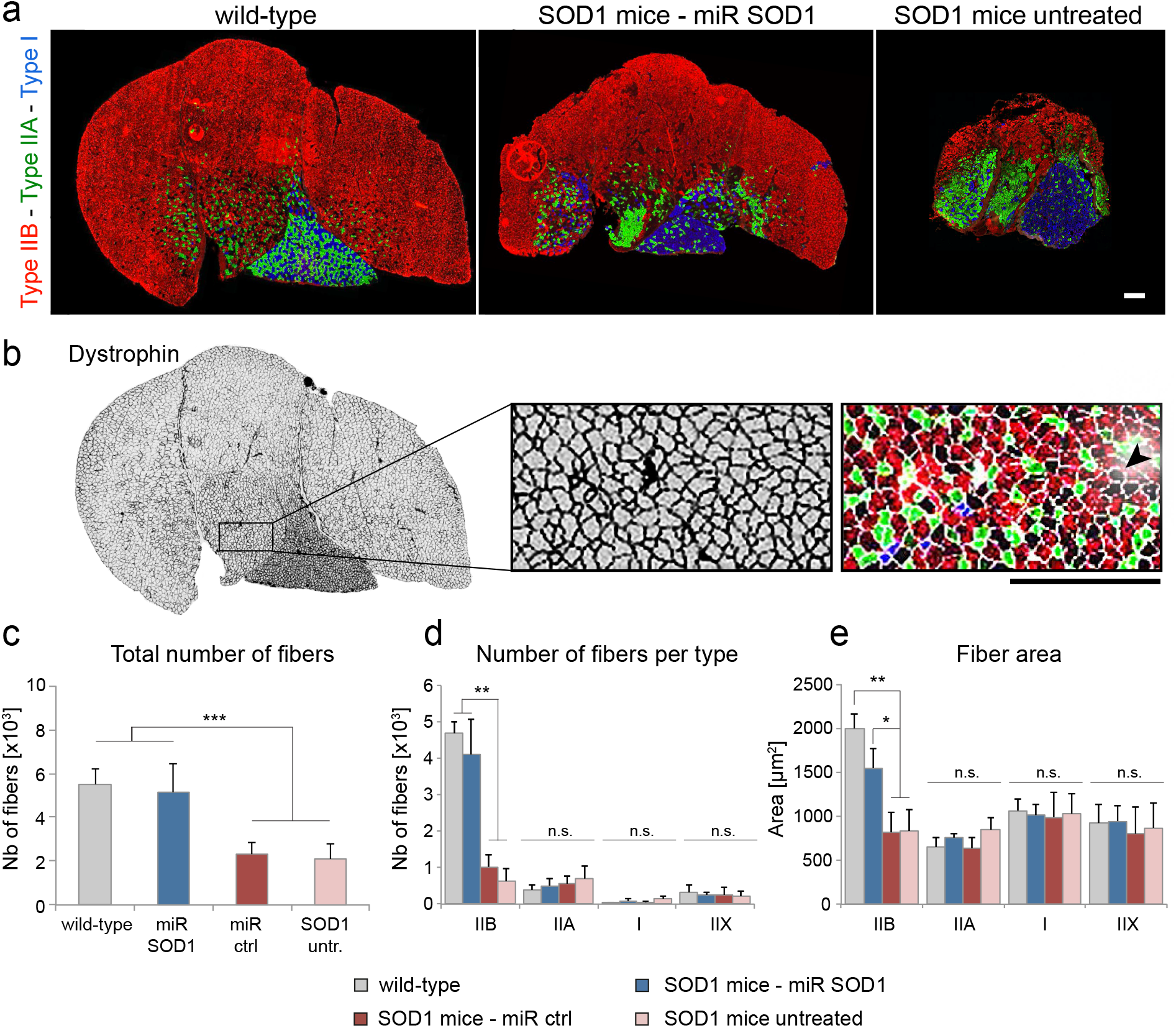
AAV miR SOD1 treatment protects against the loss of type IIb muscle fibers. (**a**) Representative images of transversal sections of the *triceps surae* muscle in WT, AAV-miR SOD1 treated and untreated ALS mice at end stage. Sections were co-immunostained for three MyHC isoforms: isoform IIb (red) shows type IIB fibers forming fast-twitch fast-fatigable motor units; isoform IIa (green) shows type IIA fibers forming fast-twitch fatigue-resistant motor units; isoform I (green) shows type I fibers forming slow-twitch motor units. Note the lack of muscle atrophy in the *SOD1^G93A^* mice treated with AAV-miR SOD1, as compared to untreated ALS mice. Scale bar: 300 μm. (**b**) Dystrophin staining (left panel) delineates individual muscle fibers, the type of which is determined according to co-immunostaining for MyHC isoform (right panel). Fibers negative for all three MyHC isoforms are defined as type IIX (arrowhead). Scale bar: 300 μm. (**c**) Quantitative analysis of the total number of fibers in the *gastrocnemius* and *plantaris* muscles at end stage. Note the significant protection against muscle fiber loss in the *SOD1^G93A^* mice treated with AAV-miR SOD. (**d**) Number of muscle fibers for each fiber type. (**e**) Mean area of individual muscle fibers, according to fiber type. Note the significant protection of the number and size of type IIb muscle fibers in AAV-miR SOD1 treated *SOD1^G93A^* mice. Data represent mean ± SD; *n* = 5 for all conditions, except WT mice (*n* = 4). Statistical analysis: one-way ANOVA with Bonferroni *post-hoc* test; n.s. not significant, \**P* < 0.05, \*\**P* < 0.01, \*\*\**P* < 0.001.

### miR SOD1 gene therapy targeted to astrocytes induces axonal sprouting and muscle fiber type grouping

The significant rescue of NMJ occupancy observed in the AAV-miR SOD1-treated group indicates that motor nerve sprouting may have occurred in the *gastrocnemius* muscle. To qualitatively assess axon sprouting events, we performed a co-staining of NMJ with α-bungarotoxin, SV2 and NFM-145, a marker for axonal neurofilaments. At day 100, terminal sprouting characterized by the presence of axons extending beyond the motor endplate was indeed observed in AAV-miR treated mice (Fig. 5a), which indicates that the treatment may have induced re-innervation of vacant endplates.

**Figure 5.**
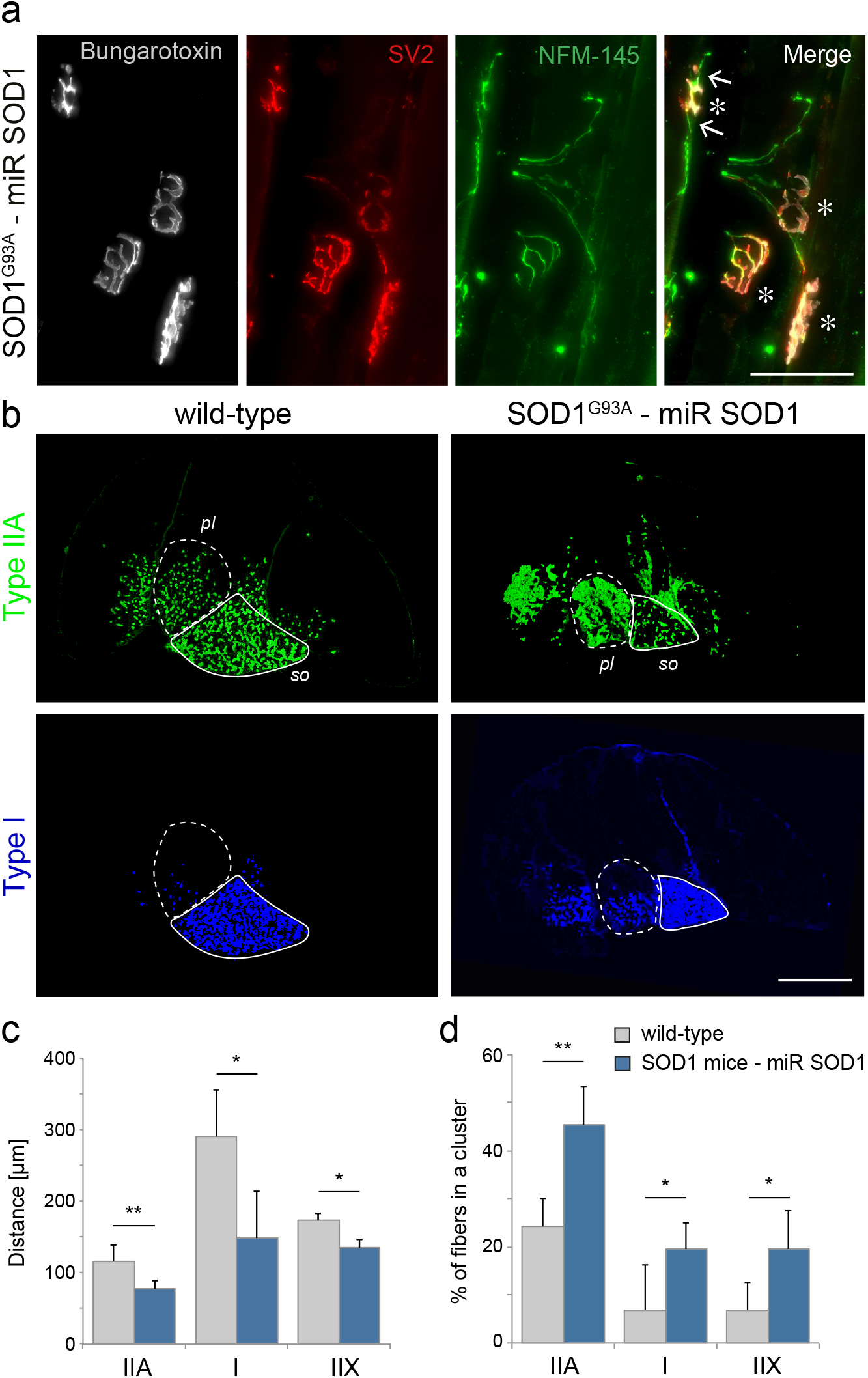
Axonal sprouting and fiber type grouping in the *triceps surae* of AAV-miR SOD1-treated *SOD1^G93A^* mice. (**a**) Representative photomicrograph of NMJ stained with α-bungarotoxin (white), SV2 (red) and NFM-145 (green) in the *gastrocnemius* muscle of an AAV-miR SOD1 treated *SOD1^G93A^* mouse at day 100. The co-localization of SV2 and α-bungarotoxin indicates complete endplate innervation in all NMJ (*). The arrows show a NFM-145-positive axon extending beyond the motor endplate, which is considered as a terminal sprouting event. Scale bar: 75μm. (**b**) Transversal sections of the *triceps surae* muscle stained for type IIa and type I MyHC to reveal the distribution of muscle fiber types in a WT and an AAV-miR SOD1-treated *SOD1^G93A^* mouse. Note the grouping of type IIA and type I muscle fibers in the treated ALS mouse (right panels), particularly evident in the *plantaris* (*pl*, dotted line) and *soleus* (*so*, solid line) muscles. Scale bar: 1 mm. (**c**) Quantification of the average minimal distance between fibers from the same type. A reduced distance is indicative of fiber type grouping. (**d**) Quantification of the percentage of fibers in direct contact with another fiber from the same type. Note that for both parameters, there is a significant increase of the clustering for all the fiber types analyzed in the AAV-miR SOD1 treated *SOD1^G93A^* mice. Data represent mean ± SD; *n* = 4 WT mice and *n* = 5 AAV-miR SOD1-treated *SOD1^G93A^* mice. Statistical analysis: two-tailed unpaired Student’s *t* test; \**P* < 0.05, \*\**P* < 0.01.

In neurogenic muscle atrophy, muscle fiber type grouping is often observed when denervation is followed by re-innervation. To assess plastic changes in the skeletal muscle, we sought to determine the level of clustering of muscle fibers by comparing WT and AAV-miR SOD1-treated ALS mice. This analysis was only possible on non-atrophied muscle tissues. While the overall number of muscle fibers was similar in both groups (see Fig. 4c), MyHC staining revealed signs of fiber-type grouping in the treated *SOD1^G93A^* mice. In Fig. 5b, this effect is particularly evident in the *plantaris* muscle, where different types of muscle fibers are intermingled. To quantify the clustering of type I, IIA and IIX muscle fibers, we measured the average distance between fibers from the same type (Fig. 5c) and the percentage of fibers that are adjacent to at least one other fiber from the same subtype (Fig. 5d). Both parameters revealed significant fiber grouping effects for all three fiber types in the *gastrocnemius* and *plantaris* muscles of AAV-miR SOD1-treated mice. As there is no loss of type IIB fibers to explain this apparent clustering in the AAV-miR SOD1-treated mice, it is likely to reflect fiber type conversion due to re-innervation of vacant endplates via the outgrowth of motor axon collaterals.

### Transcriptional signature of MN in the spinal cord of miR SOD1-treated ALS mice reveals changes in genes controlling microtubule stability

Next, we sought to explore the changes in gene expression induced by the presence of mSOD1-expressing astrocytes in the lumbar spinal cord. Similar to our previous experiment, *SOD1^G93A^* mice were ICV injected at 2 days of age either with the AAV-miR SOD1 vector or with AAV-miR ctrl. An additional group of non-injected *SOD1^G93A^* littermate mice was included in the experiment. At day 65, a transcriptomic analysis was performed on MN captured by laser microdissection in the ventral horn. This time point was selected for analysis of gene expression across conditions, as the number of surviving MN was previously found to be very similar in each group (see Fig. 2b), avoiding any confounding effects due to differences in MN survival. Whole transcriptome was analyzed by next generation sequencing, reaching on average 5.3E7 ± 1.0E7 reads per sample (mean ± SD), out of which 4.2E7 ± 0.9E7 reads where aligned to the reference mouse genome With the statistical tool DESeq2, we found 95 genes with changes in expression induced by SOD1 silencing with a false discovery rate (FDR) below 10% (Fig. 6a). Sixty-four genes were significantly upregulated and 31 genes downregulated in the AAV-miR SOD1 mice, as compared to untreated and AAV-miR ctrl injected *SOD1^G93A^* mice. For several differentially expressed genes, the changes in gene expression were confirmed by qPCR (Supplemental Table 1). A gene ontology analysis for biological processes was performed with the software GOrilla (http://cbl-gorilla.cs.technion.ac.il/), taking into account the genes with a FDR below 20% (473 genes) as compared to the total list of identified genes. Analysis showed a significant enrichment for genes involved in heme biosynthesis (*P* = 8.15E-9), chromatin organization (*P* = 8.53E-5), microtubule stability (*P* = 5.24E-4) and synaptic glutamatergic transmission (*P* = 7.49E-4).

**Figure 6:**
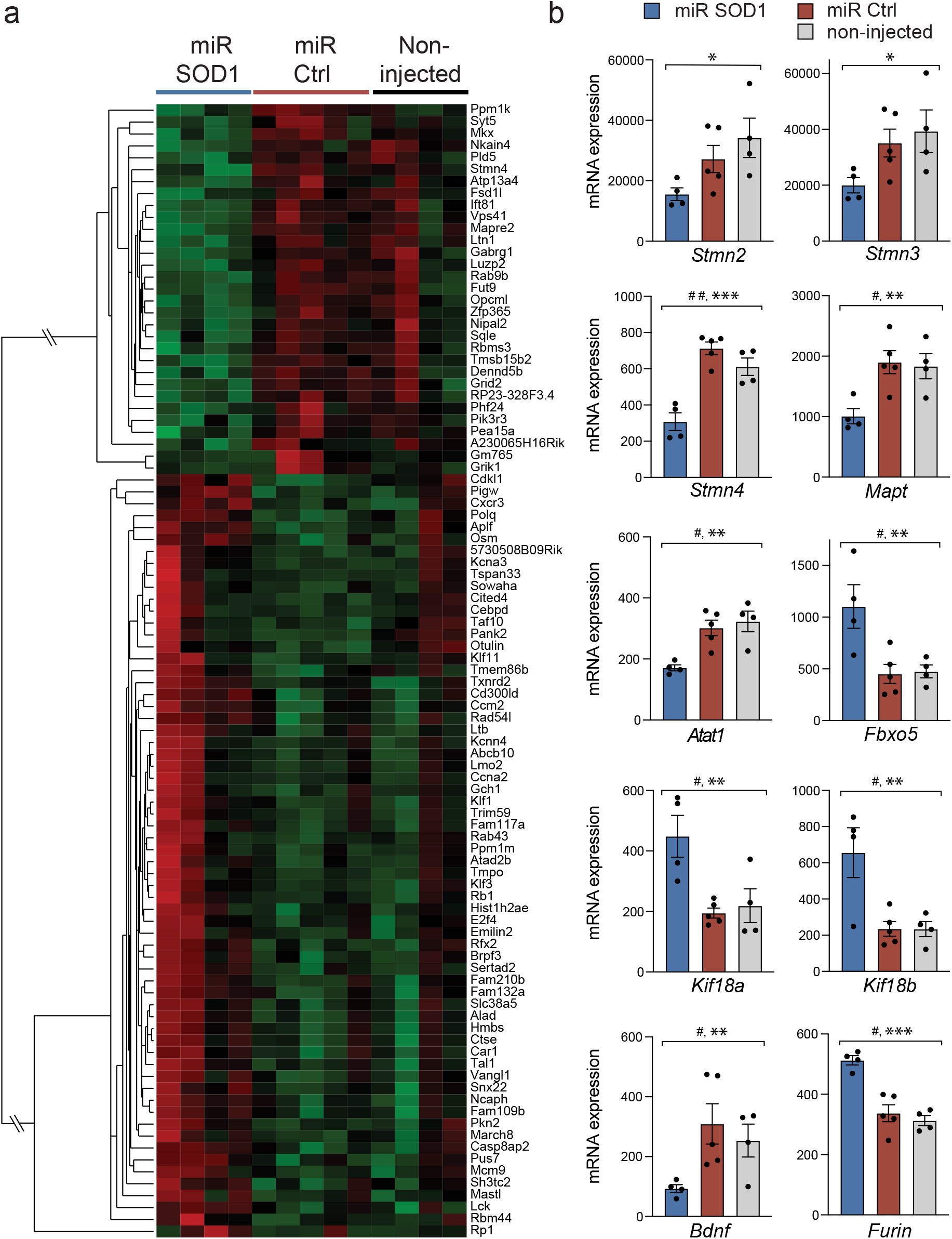
Differentially expressed genes in spinal cord motoneurons of *SOD1^G93A^* mice following AAV-miR SOD1 targeting of astrocytes. A multi-factorial statistical model with DESeq2 is used to identify 95 genes differentially expressed in MN of AAV-miR SOD1 *SOD1^G93A^* treated mice (*n* = 4) as compared to AAV-miR ctrl (*n* = 5) and non-injected *SOD1^G93A^* mice (*n* = 4). (**a**) Genes with expression changes at a false discovery rate (FDR) ≤ 10% are shown in the hierarchical clustering dendrogram. (**b**) Histogram plots showing differences in mRNA expression for individual genes either regulating microtubule stability or related to MN plasticity. Data represent mean ± SEM. Statistical analysis: FDR adjusted *P* value: # *P*adj < 0.1, ## *P*adj < 0.2; one-way ANOVA: \**P* < 0.05, \*\**P* < 0.01, \*\*\**P* < 0.001.

We sought for gene expression changes related to the observed effects of AAV-miR SOD1 on neuronal plasticity. As compared to control conditions, we found three genes of the Stathmin family to be downregulated (*Stmn2*, *Stmn3* and *Stmn4*) (Fig. 6b). Stathmins are involved in neuronal plasticity and have been shown to destabilize microtubules and inhibit their polymerization by sequestering αβ-tubulin heterodimers ^28^. The *Mapt* gene encoding the microtubule-binding protein tau was also downregulated in the AAV-miR SOD1 condition (Fig. 6b). In addition, expression of the α-tubulin-acetyl transferase gene *Atac1* was significantly downregulated, in contrast to the expression of the kinesin motor proteins Kif18a and Kif18b, which were upregulated following SOD1 silencing (Fig. 6b). Overall, these changes in gene expression showed that the AAV-miR SOD1 treatment affects microtubule dynamics in spinal cord MN, at the level of factors controlling the polymerization, stability and post-translational modifications of tubulin. Furthermore, the expression of *Fbxo5* was significantly upregulated in the treated mice (Fig. 6b). *Fbxo5* is a suppressor of Cdh1 implicated in axoneogenesis in the adult CNS ^29^. AAV-miR SOD1 treated mice showed downregulation of BDNF (Fig. 6b), a neurotrophic factor implicated in the competition between axons for making neuromuscular connections ^30^. The effects of BDNF depend on the proteolytic conversion from the pro-to the mature form of the neurotrophic factor. Indeed, the transcription of *Furin*, a key enzyme in this process, was increased in AAV-miR SOD1-injected mice (Fig. 6b).

Changes in MN activity were previously observed in models of SOD1-related ALS, shifting from hyper-to hypoexcitability in adult spinal motoneurons ^31,32^. In addition, expression of mSOD1 affects the ability of astrocytes to regulate glutamate receptor expression in MN ^33^. Here, AAV-miR SOD1 had a significant effect on the expression of several genes involved in synaptic neurotransmission. Notably, subunit-encoding genes of the ionotropic glutamate receptor family such as *Gria1* (glutamate ionotropic receptor 1, AMPA1) and *Grid2* (glutamate ionotropic receptor delta type subunit 2) were significantly downregulated (Supplemental Fig. S4a, b). Similarly, the levels of the transcripts encoding the gamma-aminobutyric acid (GABA) A receptor subunits γ1 and γ2 were significantly reduced in the AAV-miR SOD1 treated condition (Supplemental Fig. S4c, d). Genes implicated in cholinergic neurotransmission such as *ChAT* (choline acetyl transferase) and *Slc5a7* (high affinity choline transporter 1) were also downregulated (Supplemental Fig. S4e, f). Changes in the expression of genes implicated in neurotransmission may reflect MN plasticity and motor circuit homeostasis following AAV-miR SOD1 gene therapy targeting astrocytes.

## Discussion

In the *SOD1^G93A^* mouse model of ALS, we show that gene therapy to express specifically in astrocytes an artificial miR driving the selective silencing of human SOD1 leads to an improvement of the neuromuscular function during the late phase of disease. These results further support the important role of astrocytes in ALS. Indeed, WT glial-restricted progenitor cells implanted into the spinal cord of *SOD1^G93A^* rats efficiently differentiate into astrocytes and improve neuromuscular function ^34^. Similar results were obtained by transplanting astrocytes derived from healthy human iPSC into the spinal cord of *SOD1^G93A^* mice ^35^. Conversely, the transplantation of astrocytes expressing mSOD1 in WT mice leads to MN dysfunction, with negative effects on the neuromuscular function ^36^.

Our results indicate that AAV-miR SOD1 gene therapy targeting astrocytes enhances the propensity of MN to form new synapses on motor end plates. Astrocytes are known to release molecules that modulate synaptic activity and also control the formation, stabilization and elimination of synapses (reviewed in ^37^). Astrocytes can secrete matricellular proteins such as thrombospondin (TSB), secreted protein acidic and rich in cysteine like1 (SPARCL1), and SPARC (an antagonist of SPARCL1 also known as osteonectin), which have been shown to regulate synaptogenesis ^38–40^. These mechanisms are typically mediated by astrocytes in close contact with synaptic connections. However, it is unclear how the astrocyte-MN crosstalk in the spinal cord may regulate the distal formation of NMJ. Factors secreted by astrocytes and Schwann cells can mediate synapse formation in spinal MN cultures ^41^. In *Drosophila*, glial cells present at the NMJ contribute to the remodeling of the synaptic connections by removing presynaptic debris and immature boutons ^42^. In vertebrates, it is mainly terminal Schwann cells which have been reported to control the formation, maintenance, repair and pruning of the synapse at the NMJ ^43^. In *SOD1^G93A^* mice, Schwann cells express Sema3a, a chemorepellent inhibiting axonal outgrowth at the NMJ of type IIB/IIX muscle fibers, which are the first fibers to lose innervation in ALS ^44^. Similarly, Sema3a is expressed by a subpopulation of astrocytes during development to control axonal guidance in alpha-MN ^45^.

AAV-miR SOD1 treatment enhances limb strength in the grip test, which is coherent with the observed protection of type IIB muscle fibers in the *triceps surae*, and the significant protection of MN with high MMP9 expression. These neurons may constitute a pool of fast-fatigable MN protected from neurodegeneration. This effect might as well reflect a gain of MMP9 expression in the remaining pool of MN, similar to what has been observed in another transgenic mouse model of ALS following suppression of the cytosolic TDP-43 transgene ^46^. It remains however unclear whether the expression of MMP9 is necessary for NMJ re-innervation. In non-diseased conditions, NMJ re-innervation following injury has been found to occur more rapidly in fast-fatigable than in slow MN ^47^. In rodent models of ALS however, sprouting events appear to mainly occur in fatigue-resistant and slow MN, whereas there is so far no evidence for similar effects in the fast-fatigable pool ^48,49^. Here, the remarkable protection of type IIB fibers indicates that gene therapy targeting astrocytes in the *SOD1^G93A^* mouse model of ALS may enhance re-innervation even in fast muscles. Fiber type grouping is observed in patients suffering from spinal and bulbar muscular atrophy, but to a lesser extent in ALS patients ^50,51^. In contrast to other neuromuscular disorders, it is therefore possible that either the ability of MN to remodel neuromuscular synaptic connections is impaired in ALS, or that disease causes extensive degeneration before muscle fiber grouping can be observed. The observed increase in clustering of type I and type IIA muscle fibers shows that the AAV-miR SOD1 treatment enhances the ability of the surviving MN to make new functional connections with the muscle towards disease end stage, and that this effect is likely to involve all types of MN in the spinal cord. Similarly, treatment of *SOD1^G93A^* ALS mice with antisense oligonucleotides (ASO) targeting SOD1 also induces a gain in CMAP amplitude at a later stage of the disease, indicating possible rescue effects also with this mode of treatment ^52^. However, it is unclear to which extent the ASO treatment targets mSOD1 expression in astrocytes.

The analysis of gene pathways in MN highlights changes in the expression of genes regulating microtubule dynamics following AAV-miR SOD1 treatment. In particular, genes of the Stathmin family are consistently downregulated, which may reflect increased microtubule stability. Previous studies have shown in a mouse model of spinal muscular atrophy that decreased expression of stathmins ameliorates neuromuscular defects ^53^. In ALS, stathmins have also been implicated in the loss of microtubules leading to Golgi fragmentation in mutant SOD1 MN and to other defects in ALS notably through its action on Stat3 (signal transducer and activator of transcription 3) ^54–56^. More recently, TDP43 has been found to be a transcriptional repressor of Stmn2, further highlighting the role of stathmin regulation in ALS ^57,58^. We found that the treatment affects the expression of other microtubule-associated proteins in MN. There is a significant downregulation of the *Mapt* gene, which has previously been associated to the risk of developing sALS ^59,60^. Here, changes in the expression level of the microtubule-binding tau protein may affect cytoskeleton dynamics, possibly via its role in stabilizing long labile microtubule domains ^61,62^. The upregulation of the kinesin-8 family members *Kif18a* and *Kif18b*, which have been reported to have microtubule depolymerizing activity may further contribute to controlling axon length ^63^. Remarkably, transcriptomic analysis also reveals changes related to the post-translational modifications of tubulin. The transcriptional downregulation of *Atat1* (alpha tubulin acetyltransferase 1) may reflect changes in the level of this enzyme controlling α-tubulin acetylation and thereby axonal branching ^64^. *Atat1* also regulates microtubule stability independently form its activity on tubulin acetylation ^65^ Remarkably, the observed downregulation of BDNF transcription is concomitant with an upregulation of the proteolytic enzyme Furin, which may facilitate the activity-dependent stabilization of NMJ via the processing of pro-BDNF ^30,66^.

Overall, the use of AAV-based therapy targeting astrocytes as a platform to silence mSOD1 in the motor system of *SOD1^G93A^* ALS mice shows major effects on the ability on the MN to maintain functional connections with muscle fibers. These results further emphasize the importance of astrocytic cells when designing gene therapies to slow down ALS progression.

## Materials and Methods

### Animals and vector administration

All animal works were performed in accordance with the Swiss legislation and the European Community Council directive (86/609/EEC) for the care and use of laboratory animals. B6.Cg-Tg(SOD1**\*** G93A)dl1Gur/J mice (The Jackson Laboratory, Bar Harbor, USA) were mated with C57BL/6J females (Charles River Laboratories, Bois des Oncins, France). Newborn pups were genotyped at birth by polymerase chain reaction (PCR) against human SOD1. ICV injections were performed on 2-days-old pups as previously described ^26^. AAV vector suspensions were diluted in a physiologic solution of sodium chloride and mixed with 0.1% Fast Green FCF (Sigma-Aldrich) to visualize spread of the vector suspension throughout the ventricles. Three microliters of viral suspension were injected into the left lateral ventricle, using a 29G insulin syringe (B. Braun, Hessen, Germany).

### Viral vector production

The engineering and production of the AAV vectors were performed as previously described ^25^. Briefly, the following microRNA sequences: miR SOD1: 5’-ATT ACT TTC CTT CTG CTC GAA-3’ and miR ctrl: 5’-AAA TGT ACT GCG CGT GGA GAC-3’ were introduced into the pre-microRNA backbone of murine miR-155**’** and further cloned under the minimal GFAP promoter gfaABC_1_D into the pAAV-MCS:gfp plasmid expression cassettes. For production of recombinant AAV9 particles, shuttle plasmids were co-transfected with the pDF9 helper plasmid into HEK293-AAV cells (Agilent Technologies, Santa Clara, USA). Cells were lysed 72h following transfection. Viral particles were sequentially purified on iodixanol (Axis-shield, Dundee, United Kingdom) and ion-exchange affinity columns (GE Healthcare, Italy). Viral genomic copies were measured by TaqMan quantitative PCR (Invitrogen) using primers recognizing the human β-globin intron. The AAV9 vectors were injected at a titer of 1.4E14 viral genomes (VG)/mL for the behavioral study and at a dose of 2.9E14 VG/mL for the MN transcriptome analysis.

### Behavioral testing and electromyography

Cohorts used for behavioral experiments were litter-matched. Evoked CMAP amplitude in the *triceps surae* was evaluated using the electromyographic apparatus (AD Instruments, Oxford, UK) as described previously ^25^. For muscle strength measurements, each mouse was hold by the tail while lifting metal grids with defined weights of 20 g, 30 g and 40 g. The maximal duration of the test was set at 30 s. Two successive trials were performed with each grid. The inter-trial interval was set at 30 s. For each grid, the score was calculated by multiplying the best time performance by the weight of the grid. For each mouse, the total score was defined as the sum of the scores obtained for each of the three grids. For the rotarod test, the mouse was placed on an accelerating rod, at a speed constantly increasing from 4 to 40 rpm, during a maximal period of 300 s. The performance was measured as the time during which the mouse was able to maintain itself on the rotating rod. For *SOD1^G93A^* mice, the end stage of the disease was defined as the time at which the animal could no longer right itself within 20 s after being placed on its flank.

### Histological analysis

Mice were sacrificed at the mentioned age or at end stage of the disease by intra-peritoneal injection of pentobarbital (Streuli Pharma, Uznach, Switzerland). Mice were transcardially perfused with PBS and one *gastrocnemius* muscle was directly embedded in Cryomatrix (Thermo Fisher Scientific), frozen on dry ice and kept at **-**80**°**C for muscle fiber analysis. Animals were further perfused with paraformaldehyde 4% (PFA, Karl Roth). The second *gastrocnemius* muscle and spinal cord were post-fixed in PFA 4% for 20 min and 90 min, respectively, before being transferred to a 25% sucrose solution.

Twenty-five μm thick sections of the lumbar spinal cord were cut on a cryostat and conserved free floating in PBS-azide solution. Twelve μm thick transversal sections of unfixed *triceps surae* muscle and 20 μm thick longitudinal sections of fixed *gastrocnemius* muscle were cut on a cryostat directly on glass slides.

For immunostaining, sections were incubated in a 0.15% Triton X-100, 2% bovine serum albumin (BSA), 3% normal horse serum blocking solution for 1 h at room temperature (RT). Sections were then incubated for 24 h at 4**°**C with primary antibodies diluted in blocking solution. Sections were washed and incubated with secondary antibody diluted in blocking solution for 1 hr at RT. Avidin–biotin/3,3′-diaminobenzidine method (Vectors Laboratories Inc. Burlingame, USA) was applied to reveal goat anti-ChAT (1:500, Chemicon Millipore, Billerica, USA), goat anti-MMP9 (1:1000, Sigma-Aldrich) and rabbit anti-IBA1 (1:2000, Wako Pure Chemical Industries, Osaka, Japan) immunostainings. Secondary antibodies were: biotinylated horse anti-goat IgG or biotinylated goat anti-rabbit IgG (1:200, Vector BA). Forty-eight h incubation of the primary antibody and nickel ammonium sulfate enhancement was used for MMP9 staining.

Other antibodies were: rabbit anti-GFAP (DakoCytomation, Glostrup, Denmark), mouse anti-SV2 (1:40, Developmental Studies Hybridoma Bank (DSHB), University of Iowa, Iowa City, USA), and rabbit anti-NFM-145 (1:500, Chemicon Millipore), goat anti-rabbit Cy3, donkey anti-mouse Cy3, goat anti-rabbit alexa 488 (1:500, Jackson ImmunoResearch Laboratories, West Grove, USA) and tetramethylrhodamine α-bungarotoxin (1:500, Invitrogen). For muscle fiber type identification, transversal sections of *triceps surae* were stained with a cocktail of antibodies from DSHB: mouse anti-MyHC I (BA-D5, 1:500), mouse anti-MyHC IIa (SC-71, 1:500), mouse anti-MyHC IIb (BF-F3, 1:100), and rabbit anti-dystrophin (1:200, Abcam). The secondary antibodies used were: goat anti-mouse IgG2b-alexa 647 (1:500), goat anti-mouse IgG1-alexa 488 (1:500), goat anti-mouse IgM-AMCA (1:200) and goat anti-rabbit cy3 (1:500) (Jackson ImmunoResearch Laboratories).

### Quantification

For cell counts in the spinal cord, one every ten sections were stained and counted. ChAT and MMP9-positive MN were manually counted in the ventral horn of the lumbar spinal cord section using an Olympus AX70 microscope (Olympus Corporation, Japan). Microglial activation was evaluated by manually counting Iba1-positive microglial cells. Astrocytic activation was assessed in the ventral horn of GFAP-stained lumbar sections. GFAP-positive total area was determined with ImageJ using percentile thresholding. For assessments of Iba1 and GFAP activation, five sections of the spinal cord representing ten ventral horns per animal were used. Pictures were taken with a 20X objective on a Leica DM5500 microscope (Leica, Wetzlar, Germany).

Neuromuscular innervation was quantified on 20 μm-deep z-stack pictures of at least 3 fields of view per *gastrocnemius*. Sixty endplates identified using α-bungarotoxin staining were counted per muscle. Endplates were categorized as denervated, completely or partial innervated, according to the co-staining with the SV2 marker. Pictures were taken with a 20X objective on a Leica DM5500 microscope.

For muscle pattern analysis, pictures of the entire muscle section were taken with a 20X objective on an Olympus slide scanner VS120-L100. Pictures were post processed with a home-made Fiji macro and analyzed with MATLAB. For each channel, a minimal gray intensity threshold was set using sections stained with the secondary antibodies alone. For a given channel, values above the threshold were considered as positive and the muscle fiber type determined accordingly. Muscle fibers with value below the threshold for all three channels were categorized as type IIx. Objects smaller than 150 μm^2^ or bigger than 5000 μm^2^ were excluded from the analyses.

Clustering analysis were done as follows: inter-fiber distances for fibers form the same subtype were determined according to their center of gravity. As each fiber is on average in contact with six other fibers, the median of the six shortest distances for each fiber was calculated. Median values were then averaged across all fibers from the same subtype. A second analysis was performed to determine the percentage of fibers in contact with another fiber from the same subtype. Briefly, the mean radius of a muscle fiber type was calculated based on the measured area of each fiber, and assuming that the fiber has a circular shape. If the distance between the centers of two fibers from the same subtype was longer than twice their radius plus two standard deviations, the fibers were categorized as not in contact with each other.

### Laser-capture microdissection of spinal cord MN, RNA sequencing and analysis

Mice were sacrificed at 65 days by intra-peritoneal injection of pentobarbital (Streuli Pharma, Uznach, Switzerland). The lumbar part of the spinal cord with the vertebrae was directly embedded in OCT and frozen at −80°C. Fourteen μm thick sections were cut on a cryostat and stained with a 1% cresyl violet solution to reveal MN in lumbar spinal cord sections and guide laser-capture microdissection (PALM microscope, 20X, Zeiss). Total RNA was extracted from a pool of 500 MN per mouse with the microRNeasy kit according to manufacturer’s instruction (Qiagen). RNA was quantified with a Qubit (fluorimeter from Life Technologies) and RNA integrity confirmed with a Bioanalyzer (Agilent Technologies). The SMARTer™ Ultra Low RNA kit from Clontech was used for reverse transcription and cDNA amplification according to manufacturer’s instructions, starting with 5 to 6 ng of total RNA as input. 200 pg of cDNA were used for library preparation using the Nextera XT kit from Illumina. Library molarity and quality was assessed with the Qubit and Tapestation using a DNA High sensitivity chip (Agilent Technologies). Pools of 6 libraries were loaded for clustering on Single-read Illumina Flow cells. Reads of 50 bases were generated on the Illumina HiSeq 2500 and 4000 sequencers. One sample was removed from the analysis because of the poor quality of the extracted RNA, another sample was removed because of low sequencing depth and three other samples were removed because histological examination showed only low GFP expression levels.

The purity-filtered reads were adapters and quality trimmed with Cutadapt (v. 1.8, ^67^). Reads matching to ribosomal RNA sequences were removed with fastq_screen (v. 0.9.3). Remaining reads were further filtered for low complexity with reaper (v. 15-065, ^68^). Reads were aligned against *Mus musculus.GRCm38.86* genome using STAR (v. 2.5.2b, ^69^). The number of read counts per gene locus was summarized with htseq-count (v. 0.6.1, ^70^) using *Mus musculus.GRCm38.86* gene annotation. Quality of the RNA-seq data alignment was assessed using RSeQC (v. 2.3.7, ^71^). Reads were also aligned to the *Mus musculus.GRCm38.86* transcriptome using STAR (v. 2.5.2b, ^69^) and the estimation of the isoform abundance was computed using RSEM (v. 1.2.31, [19]).

Statistical analysis was performed in R (R version 3.4.0) on 18,371 protein coding genes with at least one read count in at least one sample. A multi-factorial statistical model including the effects of (1) SOD1 silencing, (2) AAV transduction (GFP) and (3) Batch was used with DESeq2 to identify 95 miR SOD1 affected genes with a false discovery rate (FDR) ≤ 10% ^73^. Prior to heatmap visualization, expression counts data were transformed with *regularized logarithm* (rlog), then batch-corrected. A gene ontology analysis for biological processes was performed with the software GOrilla (http://cbl-gorilla.cs.technion.ac.il/) comparing the list of differentially expressed genes (FDR ≤ 20%) to the total list of identified genes. The set of RNA sequencing data is available through GEO with ID GSE148901.

### Quantitative PCR

cDNA from the SMARTer™ Ultra Low RNA kit from Clontech used for reverse transcription of the RNA obtained by laser micro-dissection were utilized to perform SYBR™ Green qPCR relative quantification. The primer pairs for specific genes were purchased from Qiagen (QuantiTect Primer Assay, listed in Table 1). Relative quantification was calculated after normalization to the reference genes X-prolyl aminopeptidase (Xpnpep1) and casein kinase 2 (Csnk2b).

### Statistical analysis

Data are presented as mean ± standard error of the mean (SEM) or mean ± standard deviation (SD) as stated in the figure legend. For most data sets, one-way ANOVA and Bonferroni *post hoc* tests were performed using MATLAB. For electromyographic measurements (CMAP), as well as performance in the grid and rotarod tests, data were analyzed using a two-way (group x time) repeated-measures ANOVA followed by a Newman-Keuls *post hoc* test using the Prism 8 software. Analyses of muscle fiber clustering comparing wild-type and treated mice was performed using two-tailed unpaired *t* test. For all statistical analyses, the α level of significance was set a 0.05.

## Acknowledgments

The Authors would like to thank Philippe Colin and Christel Voize for their technical support. We also thank Aline Aebi, Fabienne Pidoux and Viviane Padrun for the technical support and production of viral vectors used in the present study; Mylène Docquier and the Institute of Genetics and Genomics of Geneva (iGE3; http://www.ige3.unige.ch/genomics-platform.php) for high-throughput DNA sequencing. This work was supported by Swiss National Science Foundation grant (310030L_156460/1 to B.L.S., P.A., F.P. and C.Ra.) and by ERANET E-Rare FaSMALS (Grant 3ER30_160673, to B.L.S. and C.Ra.). This research work was also in part supported by a generous donation from Drs Paul and Christa Plichta to honor Mr Boris Canessa.

The Authors declare that there is no conflict of interest.

## Authors contributions

Conceptualization, C.Ro., N.B.M., C.Ra., F.E.P. and B.L.S.; Methodology, C.Ro., F.E.P., S.P. and B.L.S.; Investigation, C.Ro., N.B.M., F.E.P. and S.P.; Writing – Original Draft, C.Ro., N.B.M. and B.L.S.; Writing – Review & Editing, C.Ra.; Funding Acquisition, C.Ra., F.E.P., P.A. and B.L.S.; Supervision, B.L.S.

