## Supplemental information for "Expression of a miRNA targeting mutated SOD1 in astrocytes induces motoneuron plasticity and improves neuromuscular function in ALS mice"

Figure S1

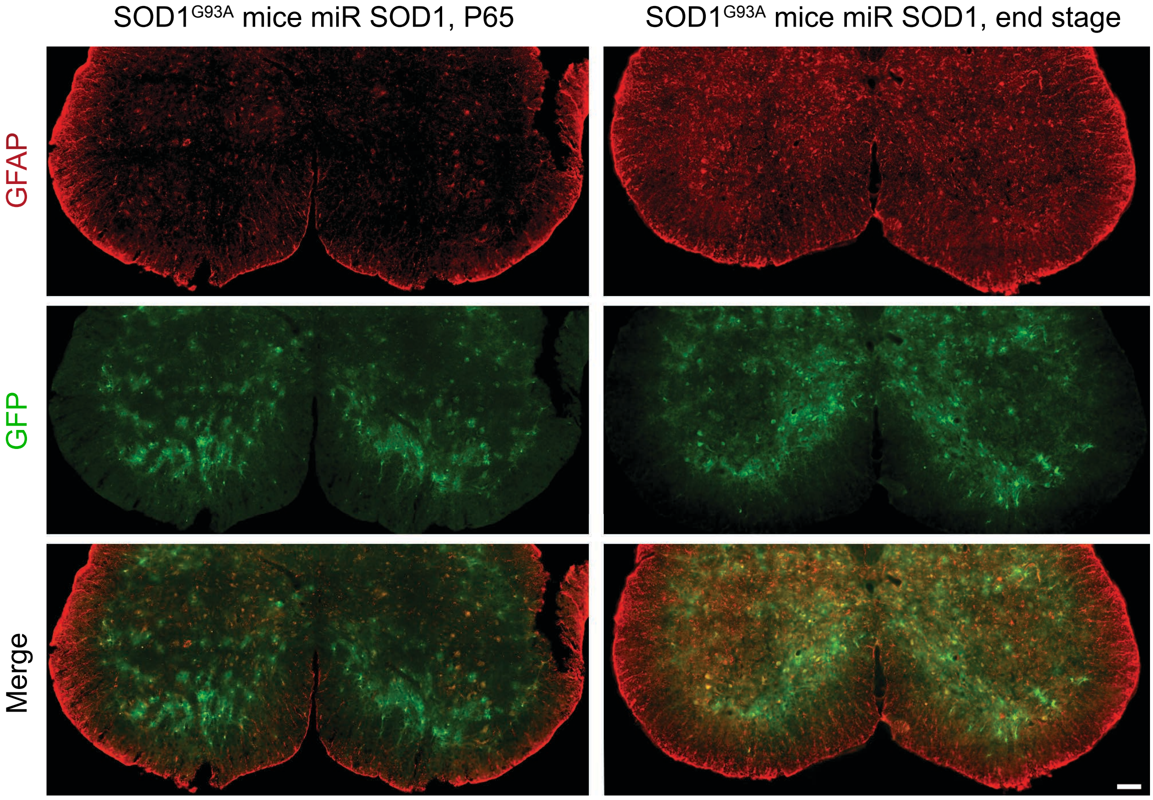


**Figure S1. GFP expression in the spinal cord over time.**

Representative pictures of spinal lumbar section stained for GFAP and GFP at day 65 and end stage. Note that GFP expression driven by the gfaABC$_1$D promoter is observed in cells with an astrocytic morphology, with a similar level at day 65 and end stage. Scale bar: 50 µm.

Figure S2

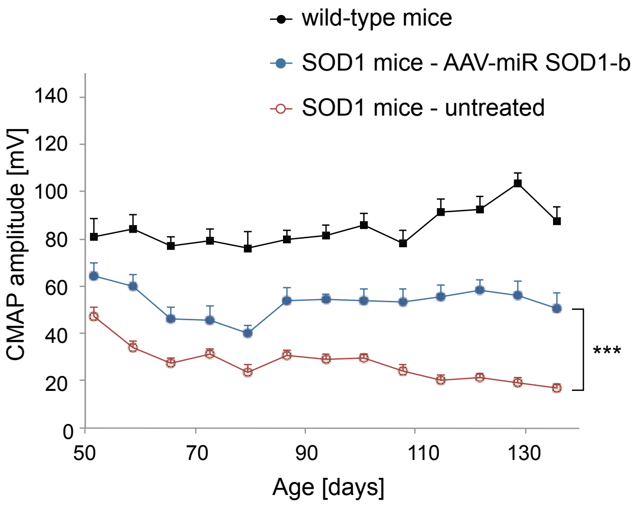


**Figure S2. Therapeutic efficacy of the AAV-miR SOD1 treatment with a second microRNA sequence targeting human SOD1.**

Amplitude of the evoked compound muscle action potential (CMAP) recorded in the *triceps surae* of non-treated wild-type and *SOD1*$^{G93A}$ mice, as compared to *SOD1*$^{G93A}$ mice injected with the AAV-miR SOD1-b vector. Note the significant rescue of CMAP values in the AAV-miR SOD1-b treated group, from day 90 on. Data represent mean ± SEM. n = 10 mice per group.

### Figure S3

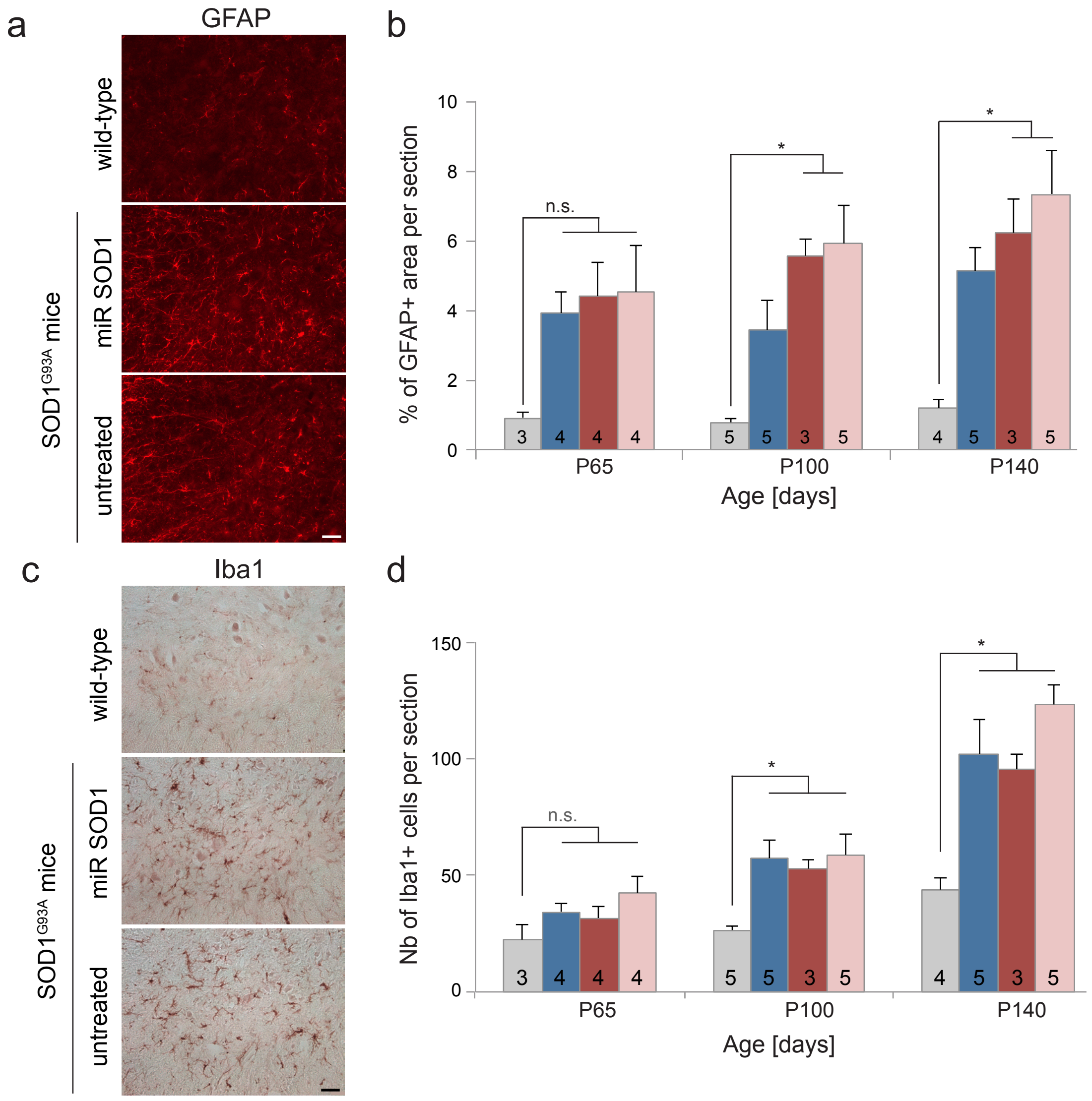


**Figure S3. AAV-miR SOD1 gene therapy targeting astrocytes does not affect astrocytic and microglial activation.**

(**a**) Representative images of GFAP immunostaining in WT and *SOD1*$^{G93A}$ mice at day 140 showing astrocytic activation in the lumbar spinal cord. (**b**) Astrogliosis is quantified by measuring the percentage of GFAP+ area in the lumbar ventral horn. Compared to WT mice, GFAP immunoreactivity is significantly increased in *SOD1*$^{G93A}$ mice at days 100 and 140. There is no significant effect of the AAV-miR SOD1 treatment. (**c**) Representative images of activated microglia immunoreactive for Iba1 in the lumbar ventral horn at day 140. (**d**) Quantification of the number of Iba1-positive activated microglial cells in the lumbar ventral horns. Compared to WT mice, a significant microglial activation is observed in *SOD1*$^{G93A}$ mice at days 100 and 140. There is no significant effect of the AAV-miR SOD1 treatment. Data represent mean ± SEM. For each graph, the numbers of replicates per group are indicated in each bar. Statistical analysis: one-way ANOVA with Bonferroni *post-hoc* test; n.s. not significant, *$P < 0.05$. Scale bars: 50 µm.

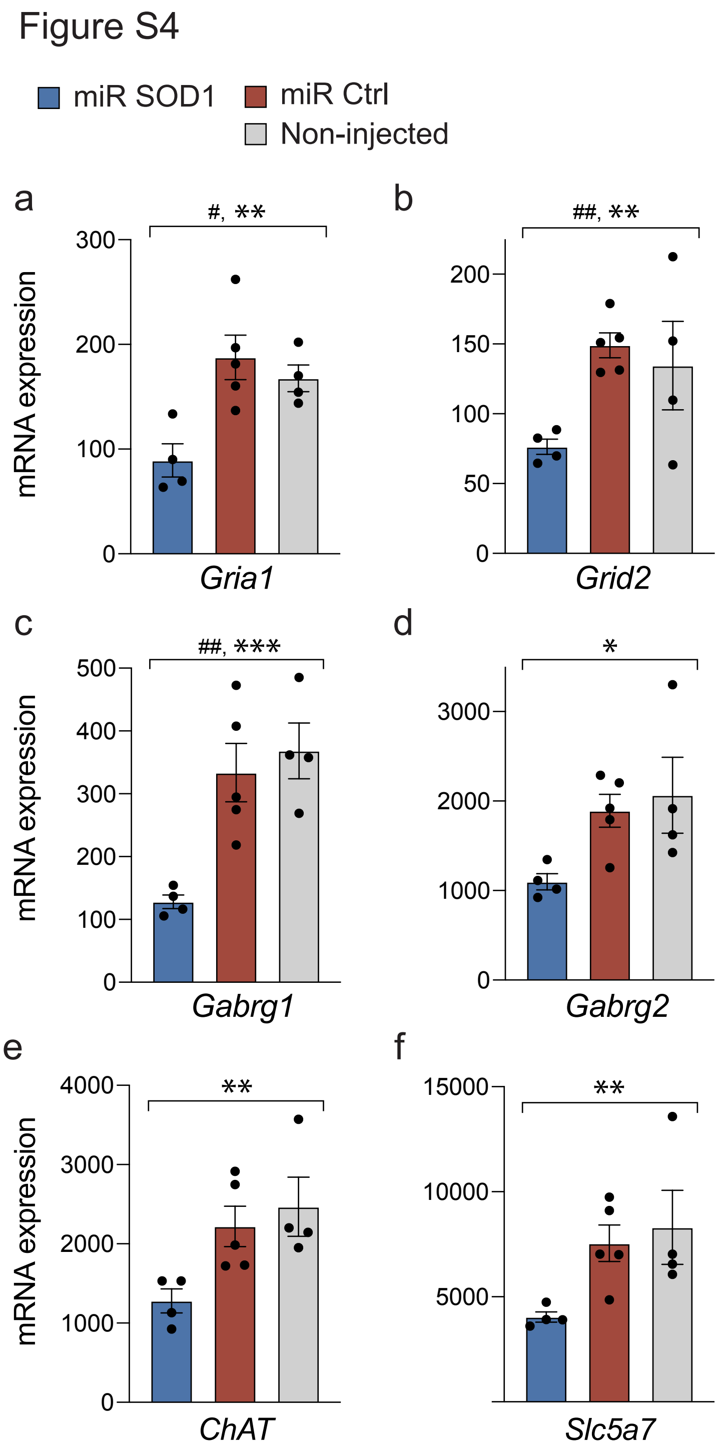


**Figure S4. Analysis of the abundance of mRNAs related to genes implicated in neurotransmission in spinal cord motoneurons of *SOD1*$^{G93A}$ mice following AAV-miR SOD1 targeting of astrocytes.**

(**a-f**) Histogram plots showing differences in mRNA expression for individual genes including glutamate receptors (*Gria1* and *Grid2*), GABA receptors (*Gabrg1* and *Gabrg2*), as well as genes involved in acetylcholine neurotransmission (*ChAT* and *Slc5a7*), in MN of AAV-miR SOD1 *SOD1*$^{G93A}$ treated mice (n = 4) as compared to AAV-miR ctrl ($n = 5$) and non-injected *SOD1*$^{G93A}$ mice ($n = 4$). Data represent mean ± SEM. Statistical analysis: False discovery rate adjusted *P* value: # *P*adj < 0.1, ## *P*adj < 0.2; one-way ANOVA: **P* < 0.05, ***P* < 0.01, ****P* < 0.001.

**Table S1: qPCR validation of differences in gene expression for a subset of genes**

| Gene symbol | Gene name | RNA seq Fold change | RNA seq FDR | qPCR Fold change | Primer set QuantiTect nb |
|---|---|---|---|---|---|
| *Abcb10* | ATP-binding cassette, sub-family B, member 10 | 4.150 | 0.034 | 1.657 | Mm_Abcb10_1_SG |
| *Alad* | aminolevulinate, delta-, dehydratase | 2.091 | 0.062 | 2.077 | Mm_Alad_1_SG |
| *Hmbs* | hydroxymethylbilane synthase | 2.816 | 0.042 | 2.569 | Mm_Hmbs_1_SG |
| *Klf1* | Krueppel-like factor 1 | 4.112 | 0.039 | 2.660 | Mm_Klf1_1_SG |
| *Lmo2* | LIM domain only 2 | 3.510 | 0.035 | 1.962 | Mm_Lmo2_1_SG |
| *Ift81* | intraflagellar transport 81 | 0.524 | 0.034 | 0.282 | Mm_Cdv1_1_SG |
| *Stmn4* | stathmin-like 4 | 0.425 | 0.011 | 0.435 | Mm_Stmn4_1_SG |
| *Trim59* | tripartite motif-containing 59 | 3.807 | 0.042 | 5.157 | Mm_Trim59_1_SG |
| *Casp8ap2* | caspase 8 associated protein 2 | 1.943 | 0.074 | 1.716 | Mm_Casp8ap2_1_SG |
| *Xpnpep1* | X-prolyl aminopeptidase 1 | 1.253 | 0.442 | - | Mm_Xpnpep1_1_SG |
| *Csnk2b* | casein kinase 2, beta polypeptide | 0.963 | 0.900 | - | Mm-Cnsk2b_1_SG |

#### Table S2: gene expression changes in motoneurons following AAV-miR SOD1 treatment

| Gene ID | Gene Symbol | log2(fold change)[a] | P value | FDR | Gene Name |
|---|---|---|---|---|---|
| ENSMUSG00000047959 | Kcna3 | 4.415 | 1.66E-06 | 0.0106 | potassium voltage-gated channel, shaker-related subfamily, member 3 |
| ENSMUSG00000022044 | Stmn4 | -1.236 | 1.79E-06 | 0.0106 | stathmin-like 4 |
| ENSMUSG00000034206 | Polq | 2.398 | 2.07E-06 | 0.0106 | polymerase (DNA directed), theta |
| ENSMUSG00000024206 | Rfx2 | 2.187 | 9.35E-06 | 0.0261 | regulatory factor X, 2 (influences HLA class II expression) |
| ENSMUSG00000034641 | Cd300ld | 1.494 | 9.58E-06 | 0.0261 | CD300 molecule like family member d |
| ENSMUSG00000027495 | Fam210b | 1.462 | 1.17E-05 | 0.0261 | family with sequence similarity 210, member B |
| ENSMUSG00000030051 | Aplf | 1.309 | 1.18E-05 | 0.0261 | aprataxin and PNKP like factor |
| ENSMUSG00000062257 | Opcml | -1.243 | 1.52E-05 | 0.0293 | opioid binding protein/cell adhesion molecule-like |
| ENSMUSG00000043463 | Rab9b | -1.377 | 2.29E-05 | 0.0337 | RAB9B, member RAS oncogene family |
| ENSMUSG00000063297 | Luzp2 | -1.261 | 2.61E-05 | 0.0337 | leucine zipper protein 2 |
| ENSMUSG00000075704 | Txnrd2 | 0.785 | 2.86E-05 | 0.0337 | thioredoxin reductase |
| ENSMUSG00000027574 | Nkain4 | -0.923 | 2.95E-05 | 0.0337 | Na+/K+ transporting ATPase interacting 4 |
| ENSMUSG00000054342 | Kcnn4 | 2.303 | 3.20E-05 | 0.0337 | K intermediate/small conductance calcium-activated channel, subfamily N, 4 |
| ENSMUSG00000036062 | Phf24 | -1.479 | 3.33E-05 | 0.0337 | PHD finger protein 24 |
| ENSMUSG00000070803 | Cited4 | 2.685 | 3.55E-05 | 0.0337 | Cbp/p300-interacting transactivator, with Glu/Asp-rich carboxy-terminal domain, 4 |
| ENSMUSG00000049800 | Sertad2 | 0.797 | 3.76E-05 | 0.0337 | SERTA domain containing 2 |
| ENSMUSG00000037826 | Ppm1k | -0.763 | 3.83E-05 | 0.0337 | protein phosphatase 1K (PP2C domain containing) |
| ENSMUSG00000031974 | Abcb10 | 2.053 | 4.23E-05 | 0.0337 | ATP-binding cassette, sub-family B (MDR/TAP), member 10 |
| ENSMUSG00000029469 | Ift81 | -0.933 | 4.25E-05 | 0.0337 | intraflagellar transport 81 |
| ENSMUSG00000004961 | Syt5 | -2.214 | 4.69E-05 | 0.0337 | synaptotagmin V |
| ENSMUSG00000001260 | Gabrg1 | -1.270 | 4.85E-05 | 0.0337 | gamma-aminobutyric acid (GABA) A receptor, subunit gamma 1 |
| ENSMUSG00000063952 | Brpf3 | 1.567 | 4.98E-05 | 0.0337 | bromodomain and PHD finger containing, 3 |
| ENSMUSG00000050549 | 5730508B09Rik | 2.405 | 5.02E-05 | 0.0337 | RIKEN cDNA 5730508B09 gene |
| ENSMUSG00000032698 | Lmo2 | 1.811 | 5.55E-05 | 0.0349 | LIM domain only 2 |
| ENSMUSG00000055373 | Fut9 | -1.079 | 5.64E-05 | 0.0349 | fucosyltransferase 9 |
| ENSMUSG00000045282 | Tmem86b | 2.159 | 6.42E-05 | 0.0381 | transmembrane protein 86B |
| ENSMUSG00000054191 | Klf1 | 2.040 | 6.87E-05 | 0.0393 | Kruppel-like factor 1 (erythroid) |
| ENSMUSG00000055214 | Pld5 | -0.985 | 7.24E-05 | 0.0399 | phospholipase D family, member 5 |
| ENSMUSG00000020990 | Cdkl1 | 1.275 | 7.62E-05 | 0.0405 | cyclin-dependent kinase-like 1 (CDC2-related kinase) |
| ENSMUSG00000034317 | Trim59 | 1.929 | 8.21E-05 | 0.0419 | tripartite motif-containing 59 |
| ENSMUSG00000032126 | Hmbs | 1.493 | 8.50E-05 | 0.0419 | hydroxymethylbilane synthase |
| ENSMUSG00000052812 | Atad2b | 1.825 | 8.84E-05 | 0.0419 | ATPase family, AAA domain containing 2B |
| ENSMUSG00000061013 | Mkx | -1.032 | 9.07E-05 | 0.0419 | mohawk homeobox |
| ENSMUSG00000004591 | Pkn2 | 1.247 | 9.22E-05 | 0.0419 | protein kinase N2 |
| ENSMUSG00000054752 | Fsd1l | -0.939 | 1.05E-04 | 0.0462 | fibronectin type III and SPRY domain containing 1-like |
| ENSMUSG00000058755 | Osm | 2.161 | 1.15E-04 | 0.0492 | oncostatin M |
| ENSMUSG00000043866 | Taf10 | 2.834 | 1.21E-04 | 0.0507 | TATA-box binding protein associated factor 10 |
| ENSMUSG00000037580 | Gch1 | 2.384 | 1.35E-04 | 0.0546 | GTP cyclohydrolase 1 |
| ENSMUSG00000031170 | Slc38a5 | 1.646 | 1.40E-04 | 0.0546 | solute carrier family 38, member 5 |
| ENSMUSG00000045140 | Pigw | 1.395 | 1.42E-04 | 0.0546 | phosphatidylinositol glycan anchor biosynthesis, class W |
| ENSMUSG00000030055 | Rab43 | 2.387 | 1.46E-04 | 0.0550 | RAB43, member RAS oncogene family |
| ENSMUSG00000027715 | Ccna2 | 2.079 | 1.50E-04 | 0.0553 | cyclin A2 |
| ENSMUSG00000050232 | Cxcr3 | 3.472 | 1.58E-04 | 0.0564 | chemokine (C-X-C motif) receptor 3 |
| ENSMUSG00000024399 | Ltb | 1.567 | 1.63E-04 | 0.0564 | lymphotoxin B |
| ENSMUSG00000044352 | Sowaha | 2.075 | 1.64E-04 | 0.0564 | sosondowah ankyrin repeat domain family member A |
| ENSMUSG00000041236 | Vps41 | -0.435 | 1.71E-04 | 0.0575 | VPS41 HOPS complex subunit |
| ENSMUSG00000090667 | Gm765 | -3.319 | 1.80E-04 | 0.0592 | predicted gene 765 |
| ENSMUSG00000028702 | Rad54l | 1.622 | 1.89E-04 | 0.0606 | RAD54 like (S. cerevisiae) |
| ENSMUSG00000026779 | Mastl | 1.718 | 1.92E-04 | 0.0606 | microtubule associated serine/threonine kinase-like |
| ENSMUSG00000020253 | Ppm1m | 1.935 | 1.98E-04 | 0.0612 | protein phosphatase 1M |
| ENSMUSG00000057541 | Pus7 | 0.975 | 2.10E-04 | 0.0621 | pseudouridylate synthase 7 |
| ENSMUSG00000039452 | Snx22 | 1.814 | 2.14E-04 | 0.0621 | sorting nexin 22 |
| ENSMUSG00000024277 | Mapre2 | -0.460 | 2.15E-04 | 0.0621 | microtubule-associated protein, RP/EB family, member 2 |
| ENSMUSG00000071637 | Cebpd | 2.381 | 2.21E-04 | 0.0621 | CCAAT/enhancer binding protein (C/EBP), delta |
| ENSMUSG00000029178 | Klf3 | 1.366 | 2.26E-04 | 0.0621 | Kruppel-like factor 3 (basic) |
| ENSMUSG00000038094 | Atp13a4 | -3.770 | 2.33E-04 | 0.0621 | ATPase type 13A4 |
| ENSMUSG00000109946 | RP23-328F3.4 | -0.884 | 2.37E-04 | 0.0621 | |
| ENSMUSG00000071424 | Grid2 | -1.196 | 2.38E-04 | 0.0621 | glutamate receptor, ionotropic, delta 2 |
| ENSMUSG00000022105 | Rb1 | 1.258 | 2.40E-04 | 0.0621 | retinoblastoma 1 |
| ENSMUSG00000028393 | Alad | 1.064 | 2.41E-04 | 0.0621 | aminolevulinate, delta-, dehydratase |
| ENSMUSG00000087075 | A230065H16Rik | -1.250 | 2.58E-04 | 0.0649 | RIKEN cDNA A230065H16 gene |
| ENSMUSG00000001763 | Tspan33 | 2.475 | 2.61E-04 | 0.0649 | tetraspanin 33 |
| ENSMUSG00000069272 | Hist1h2ae | 2.780 | 2.72E-04 | 0.0663 | histone cluster 1, H2ae |

| | | | | | |
|---|---|---|---|---|---|
| ENSMUSG00000027860 | Vangl1 | 1.358 | 2.75E-04 | 0.0663 | vang-like 1 (van gogh, Drosophila) |
| ENSMUSG00000022935 | Grik1 | -3.332 | 2.80E-04 | 0.0665 | glutamate receptor, ionotropic, kainate 1 |
| ENSMUSG00000052299 | Ltn1 | -0.520 | 2.89E-04 | 0.0676 | listerin E3 ubiquitin protein ligase 1 |
| ENSMUSG00000028717 | Tal1 | 1.388 | 2.94E-04 | 0.0676 | T cell acute lymphocytic leukemia 1 |
| ENSMUSG00000038893 | Fam117a | 1.499 | 2.98E-04 | 0.0676 | family with sequence similarity 117, member A |
| ENSMUSG00000000378 | Ccm2 | 0.614 | 3.05E-04 | 0.0676 | cerebral cavernous malformation 2 |
| ENSMUSG00000045629 | Sh3tc2 | 1.466 | 3.06E-04 | 0.0676 | SH3 domain and tetratricopeptide repeats 2 |
| ENSMUSG00000034906 | Ncaph | 1.207 | 3.18E-04 | 0.0692 | non-SMC condensin I complex, subunit H |
| ENSMUSG00000019961 | Tmpo | 1.494 | 3.47E-04 | 0.0737 | thymopoietin |
| ENSMUSG00000028698 | Pik3r3 | -1.036 | 3.48E-04 | 0.0737 | phosphatidylinositol 3 kinase, regulatory subunit, polypeptide 3 (p55) |
| ENSMUSG00000020653 | Klf11 | 1.492 | 3.56E-04 | 0.0742 | Kruppel-like factor 11 |
| ENSMUSG00000014859 | E2f4 | 1.156 | 3.64E-04 | 0.0744 | E2F transcription factor 4 |
| ENSMUSG00000025702 | March8 | 1.609 | 3.68E-04 | 0.0744 | membrane-associated ring finger (C3HC4) 8 |
| ENSMUSG00000028282 | Casp8ap2 | 0.958 | 3.71E-04 | 0.0744 | caspase 8 associated protein 2 |
| ENSMUSG00000038879 | Nipal2 | -1.418 | 3.87E-04 | 0.0766 | NIPA-like domain containing 2 |
| ENSMUSG00000037514 | Pank2 | 1.437 | 4.08E-04 | 0.0767 | pantothenate kinase 2 |
| ENSMUSG00000013698 | Pea15a | -0.664 | 4.11E-04 | 0.0767 | phosphoprotein enriched in astrocytes 15A |
| ENSMUSG00000089996 | Tmsb15b2 | -1.450 | 4.14E-04 | 0.0767 | thymosin beta 15b2 |
| ENSMUSG00000023571 | Fam132a | 1.092 | 4.17E-04 | 0.0767 | family with sequence similarity 132, member A |
| ENSMUSG00000058298 | Mcm9 | 1.877 | 4.17E-04 | 0.0767 | minichromosome maintenance 9 homologous recombination repair factor |
| ENSMUSG00000024053 | Emilin2 | 1.612 | 4.17E-04 | 0.0767 | elastin microfibril interfacer 2 |
| ENSMUSG00000070732 | Rbm44 | 3.304 | 4.29E-04 | 0.0774 | RNA binding motif protein 44 |
| ENSMUSG00000004552 | Ctse | 1.698 | 4.31E-04 | 0.0774 | cathepsin E |
| ENSMUSG00000039607 | Rbms3 | -0.698 | 4.74E-04 | 0.0841 | RNA binding motif, single stranded interacting protein |
| ENSMUSG00000027556 | Car1 | 1.659 | 4.89E-04 | 0.0858 | carbonic anhydrase 1 |
| ENSMUSG00000022351 | Sqle | -0.728 | 4.96E-04 | 0.0860 | squalene epoxidase |
| ENSMUSG00000030313 | Dennd5b | -0.763 | 5.52E-04 | 0.0946 | DENN/MADD domain containing 5B |
| ENSMUSG00000000409 | Lck | 1.507 | 5.58E-04 | 0.0946 | lymphocyte protein tyrosine kinase |
| ENSMUSG00000037855 | Zfp365 | -0.802 | 5.67E-04 | 0.0946 | zinc finger protein 365 |
| ENSMUSG00000025900 | Rp1 | 5.495 | 5.70E-04 | 0.0946 | retinitis pigmentosa 1 (human) |
| ENSMUSG00000049687 | Fam109b | 1.362 | 5.90E-04 | 0.0969 | family with sequence similarity 109, member B |
| ENSMUSG00000046034 | Otulin | 1.743 | 6.14E-04 | 0.0997 | OTU deubiquitinase with linear linkage specificity |

[a]miRSOD1 vs [miR ctrl and non-inj.]
